# Purification of native CCL7 and its functional interaction with selected chemokine receptors

**DOI:** 10.1101/771618

**Authors:** Marina V. Goncharuk, Debarati Roy, Maxim A. Dubinnyi, Kirill D. Nadezhdin, Ashish Srivastava, Mithu Baidya, Hemlata Dwivedi-Agnihotri, Alexander S. Arseniev, Arun K. Shukla

## Abstract

Chemokine receptors form a major sub-family of G protein-coupled receptors (GPCRs) and they are involved in a number of cellular and physiological processes related to our immune response and regulation. A better structural understanding of ligand-binding, activation, signaling and regulation of chemokine receptors is very important to design potentially therapeutic interventions for human disorders arising from aberrant chemokine signaling. One of the key limitations in probing the structural details of chemokine receptors is the availability of large amounts of purified, homogenous and fully functional chemokine ligands, and the commercially available products, are not affordable for in-depth structural studies. Moreover, production of uniformly isotope-labeled chemokines, for example, suitable for NMR-based structural investigation, also remains challenging. Here, we have designed a streamlined approach to express and purify the human chemokine CCL7 as well as its ^15^N-, ^15^N/^13^C-, ^2^H/^15^N/^13^C-isotope-labeled derivatives, at milligram levels using *E. coli* expression system. Purified CCL7 not only maintains a well-folded three-dimensional structure as analyzed using circular dichroism and ^1^H/^15^N NMR but it also induces coupling of heterotrimeric G-proteins and β-arrestins for selected chemokine receptors in cellular system. Our strategy presented here may be applicable to other chemokines and therefore, provide a potentially generic and cost-effective approach to produce chemokines in large amounts for functional and structural studies.

## Introduction

G protein-coupled receptors (GPCRs) are responsible for recognizing a broad range of ligands at the cell surface and transmitting the signal across the membrane for downstream signaling and functional response (Bockaert and Pin, 1999). Considering their integral role in numerous pathophysiological conditions, they are one of the most sought-after drug targets, and nearly one third of the currently prescribed medicines exert their actions through this class of receptors (Kumari et al., 2015, Sriram and Insel, 2018). One of the major sub-families of GPCRs recognizes various chemokines in our body and they are collectively grouped as chemokine receptors (Hughes and Nibbs, 2018, Griffith et al., 2014). These chemokine receptors are critically important for cellular migration among many other functions, especially in the immune system (Griffith et al., 2014).

Chemokines are small proteins with less than hundred amino acid residues and they typically have overall conserved structural features (Turner et al., 2014). An interesting feature of chemokine receptors is their ligand promiscuity where a given chemokine can interact with, and modulate several related chemokine receptors (Proudfoot, 2002). Investigating the interaction of chemokines with their receptors in terms of precise ligand-receptor contacts, receptor activation, signaling and regulation is important to understand how the receptor promiscuity and preferences are encoded. Unlike small peptide GPCR ligands which can be chemically synthesized, chemokines typically harbor several disulphide linkages and require proper cellular context for folding and attaining a functional conformation.

Although several chemokines are commercially available, using them for structural studies that require milligram amounts is not feasible. Moreover, batch-to-batch variation and lack of proper functional characterization of these products further limit their usage in structural and functional studies. A number of previous studies have documented the purification of chemokines from inclusion bodies after recombinant expression in *E. coli*, however, their proper refolding and functionality remains a concern. On the other hand, preparing large amounts of chemokines from baculovirus and mammalian expression system may also not be cost-effective. Thus, new strategies to express and purify native chemokines, preferably in soluble form without refolding step, is desirable to propel structure-function studies of chemokine-chemokine receptor complexes. Moreover, streamlined methods to express and purify isotope labeled chemokines for NMR-based structural characterization is also desirable.

Here, we have designed and optimized a streamlined protocol for expression and purification of native (i.e. untagged) chemokine CCL7 in soluble form using *E. coli*. Our approach allowed us to also produce sufficient amounts of isotope-labeled derivatives for structural studies using NMR. We observed that native CCL7 purified from *E. coli* behaves as a full agonist for chemokine receptor CXCR2 and atypical chemokine receptor ACKR2a (also referred to as decoy D6 receptor; D6R) in G-protein coupling and β-arrestin recruitment assays. The strategy described here should be applicable to other chemokines and it should facilitate structural characterization of chemokine-chemokine receptor complexes in future.

## Results

### Expression construct and purification of his6-tagged CCL7

We first designed an expression construct for his6-tagged CCL7 (referred to as his-CCL7 hereafter) in a pGEMEX1 derived vector (Goncharuk et al., 2018) (Figure 1A) and optimized expression conditions by systematic comparison of expression time, temperature and IPTG concentrations. We evaluated the soluble expression of his-CCL7 using a previously published protocol (Goncharuk et al., 2011) and identified that its expression is optimal when cultures are grown at 20°C for 48h after induction with 1mM IPTG in TB (Terrific Broth) medium. For growth in minimal salt medium, cells were grown at 27°C for 48h after induction with 0.5mM IPTG induction to obtain maximal his-CCL7 expression. We followed these conditions for expression scale-up and purified his-CCL7 using a two-step purification scheme involving Ni-NTA agarose and cation-exchange chromatography. We obtained highly pure his-CCL7 with an overall yield of about 25-30mg per liter culture (Figure 1B). In order to evaluate the conformational homogeneity of purified his-CCL7, we analyzed the purified sample on Superdex 200 Increase (10/300 GL) based size exclusion chromatography which revealed a monodisperse elution profile (Figure 1C). The purity and identity of his-CCL7 was further confirmed by LC-MS which displays a predominant peak at m/z of 10,050.4 Da and confirms the formation of two disulphide bonds (Figure 1D).

**Figure 1:**
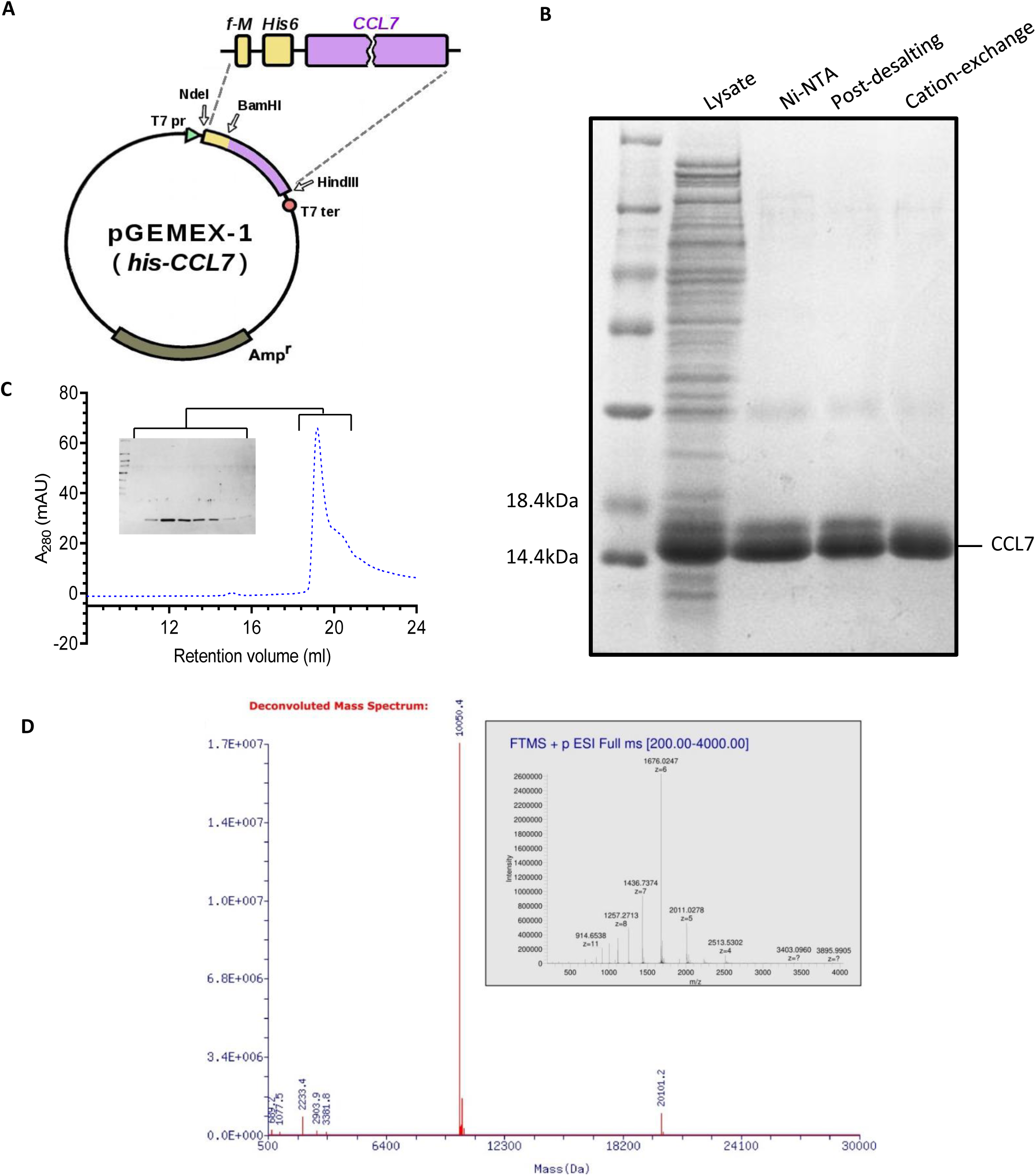
Expression and purification of his-CCL7 from *E. coli*. **A.** pGEMEX-1 derived expression vector for recombinant production of N-terminal His-tagged CCL7. The natural signal sequence of CCL7 was removed during cloning of CCL7 and a his6-tag was engineered at the N-terminus of CCL7. **B.** Purification of CCL7 using Ni-NTA, desalting and cation exchange chromatography. Samples from every step were loaded and separated using 16% Tris-Glycine SDS-PAGE followed by Coomassie staining for visualization. **C.** Size exclusion chromatography of CCL7 on Superdex 200 column revealed a monodisperse peak with an elution volume of 20mL. The inset shows the corresponding fractions analyzed by SDS-PAGE and Coomassie staining. **D.** Deconvoluted mass spectrum of his-CCL7 displays predominant m/z 10050.4 Da and confirms formation of two disulphide bonds. Inset: raw mass spectrum of his-CCL7 acquired on Thermo Scientific LTQ Orbitrap, electrospray ionization in positive mode.

### NMR analysis of ^15^N labeled his6-tagged CCL7

In order to test whether CCL7 purified using this protocol is properly folded, we confirmed its secondary structure using CD spectroscopy and NMR spectroscopy (Figure 2A-B). The observed secondary structure in CD spectroscopy agrees well with the expected range for his-CCL7 (Figure 2A). For NMR spectroscopy, we expressed and purified stable isotope labeled his-CCL7 by growing the cells in M9 minimal media containing the corresponding source of stable isotopes. Subsequently, we used unlabeled (0.3mM) and ^15^N-labeled (0.8 mM) his-CCL7 samples in H_2_O/D_2_O at pH 5.1 to acquire NMR data at 700MHz spectrometer for structural verification and sequential NMR assignment (Figure 2). We used non-uniform sampling technique (Kazimierczuk and Orekhov, 2015) to increase the resolution of 3-D ^15^N-TOCSY-HSQC (20% fill of full time domain matrix) and 3-D ^15^N-NOESY-HSQC experiment (38% fill of matrix) resulting in reduction of experimental time to 24h and 58h, respectively. For assignment, we adopted a previously reported entry for CCL7 (BMRB entry 4177) (Kim et al., 1996) as it does not have any significant difference with our construct except the Gly-Ser sequence between his6 and CCL7. We observed that all the NMR NOESY contacts support well-known spatial structure of CCL7 published previously in PDB, for example, by NMR (1BO0 and 1NCV, monomer and dimer, respectively) (Kim et al., 1996, Meunier et al., 1997) and by X-ray crystallography (4ZKC)(Counago et al., 2015). For example, the α-helix K58-L67, and β-sheets L25-R30 and V41-T45, are clearly identified by their characteristic NOE contacts (data not shown). Interestingly, we did not observe any of the eight inter-monomer NOE contacts reported previously for CCL7 (Meunier et al., 1997) suggesting that his-CCL7 purified here is primarily in a monomeric state. It is also plausible that our ^15^N-resolved 3D experiments at high resolution provide a more accurate analysis of ambiguous NOE restraints compared to earlier work using 2D NMR of unlabeled CCL7 (Meunier et al., 1997).

**Figure 2:**
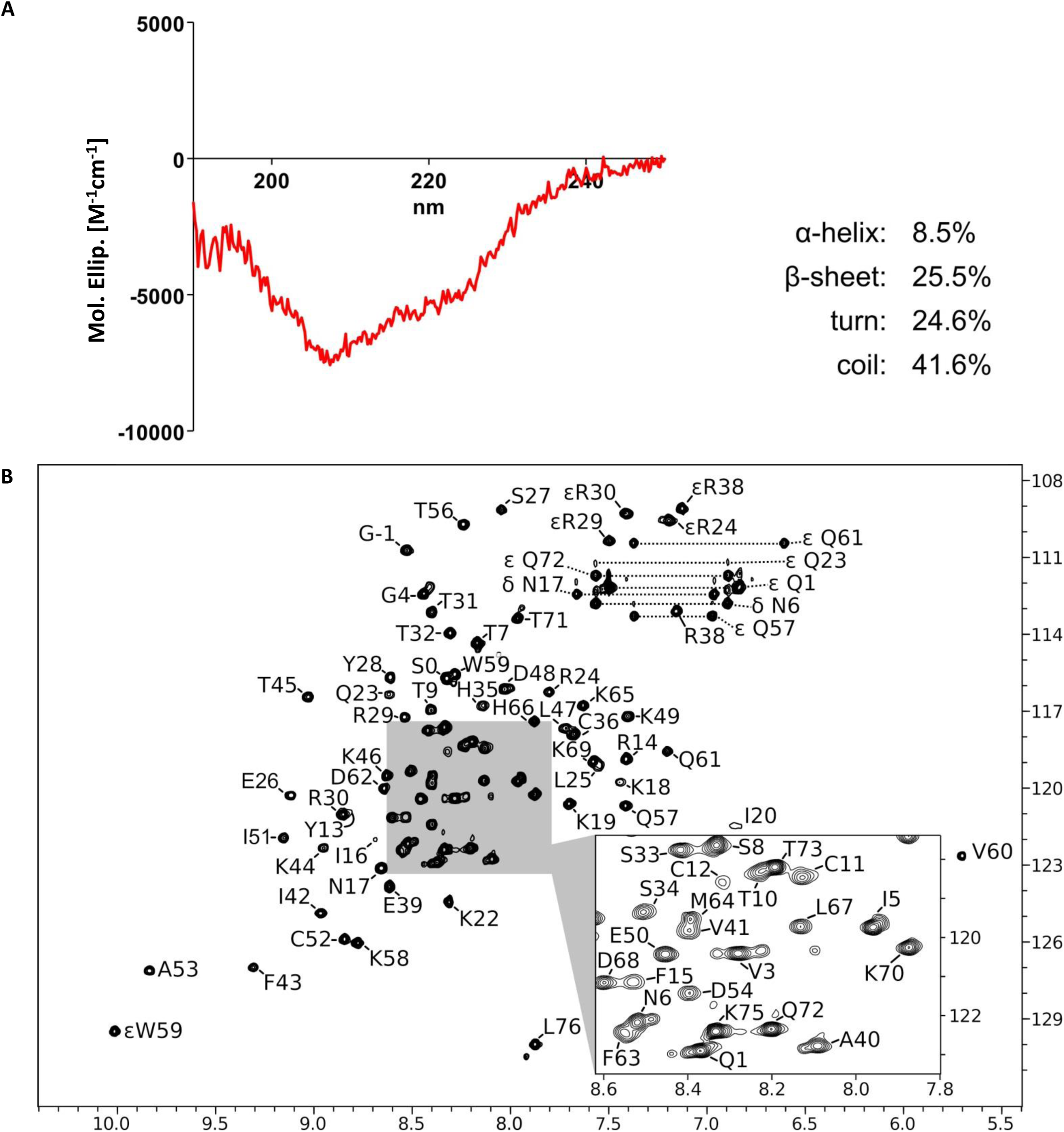
Structural characterization of his-CCL7 by CD spectroscopy and NMR. **A.** Circular dichroism (CD) spectrum of his-CCL7 acquired on J-810 spectropolarimeter (JASCO, Japan) at pH 5.8. The percentage of secondary structure elements were calculated in CONTINLL software (CDPro package) using SMP56 reference spectra. **B.** ^1^H-^15^N HSQC NMR spectrum of ^15^N-labeled his-CCL7. Backbone NH groups are denoted by one-letter amino acid notation followed by the residue number. NMR signals of N-terminal his6-tag are not observed. Non-natural N-terminal GS-linker residues are denoted as G-1 and S0 and together with the visible NH group of Q1 that three signals constitute all differences with natural CCL7 in ^1^H-^15^N HSQC as described earlier in (Kim et.al. 1996). The NMR signals of side-chain amines are prefixed by greek letters, two related resonances of NH_2_ groups (N, Q) are linked by dotted lines. Signals of εR are folded in ^15^N dimension (real ^15^N chemical shift ∼85 ppm). Central part of the spectrum is expanded for clarity. Chemical shifts of ^1^H and ^15^N are in ppm referenced to water resonance (4.7ppm). Conditions: Bruker Avance 700 MHz, 320 μL of 0.8 mM of His-CCL7 in H_2_O:D_2_O (9:1) pH 5.1, shigemi tube, temperature 30°C.

### Functional characterization of his-CCL7 in G-protein coupling assay

In order to probe the functionality of purified his-CCL7, we measured its ability to induce Gαi-coupling upon stimulation of chemokine receptor CCR2 using GloSensor assay (Kumar et al., 2017). We used carboxyl-terminal Fc-tagged CCL7 (CCL7-Fc) purified from *Sf*9 cells as a reference. Although his-CCL7 behaved as a full-agonist at CCR2 with respect to Gαi-coupling with almost an equivalent B_max_ as CCL7-Fc, its potency was approximately 100 fold less than that of CCL7-Fc (IC_50_ of ∼10nM)(Figure 3A-B).These observations suggest that the his6-tag present at the N-terminus of CCL7 potentially interferes with its ability to induce effective G-protein-coupling at CCR2. Thus, we designed an alternative strategy to generate a native CCL7 without any N-terminal or C-terminal tag.

**Figure 3:**
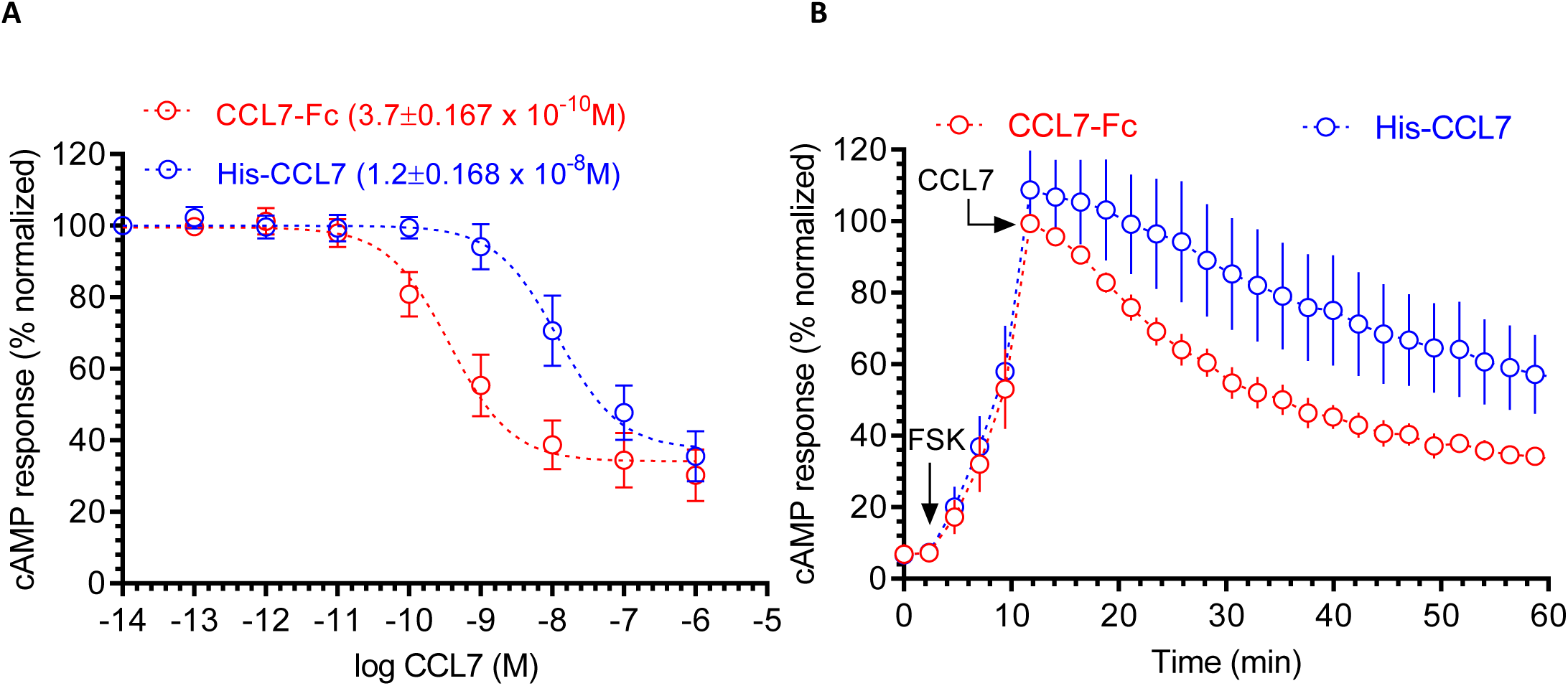
Functional characterization of his-CCL7 in GloSensor based G-protein coupling assay. **(A)** The ability of different CCL7 preparations to inhibit Forskolin-induced cAMP response downstream of human chemokine receptor CCR2 in transfected HEK-293 cells. The his-CCL7 represents N-terminal His6-tagged version while CCL7-Fc represents a C-terminal Fc-tagged version of CCL7 expressed and purified from *Sf*9 cells. Data represent average±SEM of four independent experiments carried out in duplicates and normalized with respect to lowest concentration of CCL7-Fc (treated as 100%). **(B)** Time-course analysis of CCL7 induced decrease in the cAMP level over the indicated time-period. Values recorded in the GloSensor assay at a concentration of 10nM from the experiments presented in panel A are plotted. The arrow indicates the time of CCL7 addition and the values are normalized with maximal cAMP response observed for CCL7-Fc (treated as 100%).

### Expression construct and purification of native CCL7

In order to generate a fully native CCL7 without any modification at the N- or the C-terminal, we designed an expression construct with N-terminal his6-tag followed by an enterokinase cleavage site referred to as his-EK-CCL7 (Figure 4A). As enterokinase effectively cleaves the fusion protein after lysine (or arginine), incorporation of its recognition sequence (Asp-Asp-Asp-Asp-Arg) before CCL7, allows the generation of native N-terminus in CCL7. In order to enhance the availability of the enterokinase cleavage site, a flexible linker (GSGSG) was engineered after the his6-tag. Similar to his6-tagged CCL7, we optimized the expression conditions for his-EK-CCL7 and purified it first using Ni-NTA chromatography (Figure 4B). Afterwards, we cleaved the fusion protein with the enterokinase which yielded approximately 70% cleavage of the fusion protein. Subsequent cation exchange chromatography using Resource S column efficiently separated the cleaved CCL7 and uncleaved fusion protein (Figure 4B). In order to evaluate the conformational homogeneity of purified native CCL7, we analyzed the purified sample on Superdex 200 Increase (10/300 GL) based size exclusion chromatography which revealed a monodisperse elution profile (Figure 4C).

**Figure 4:**
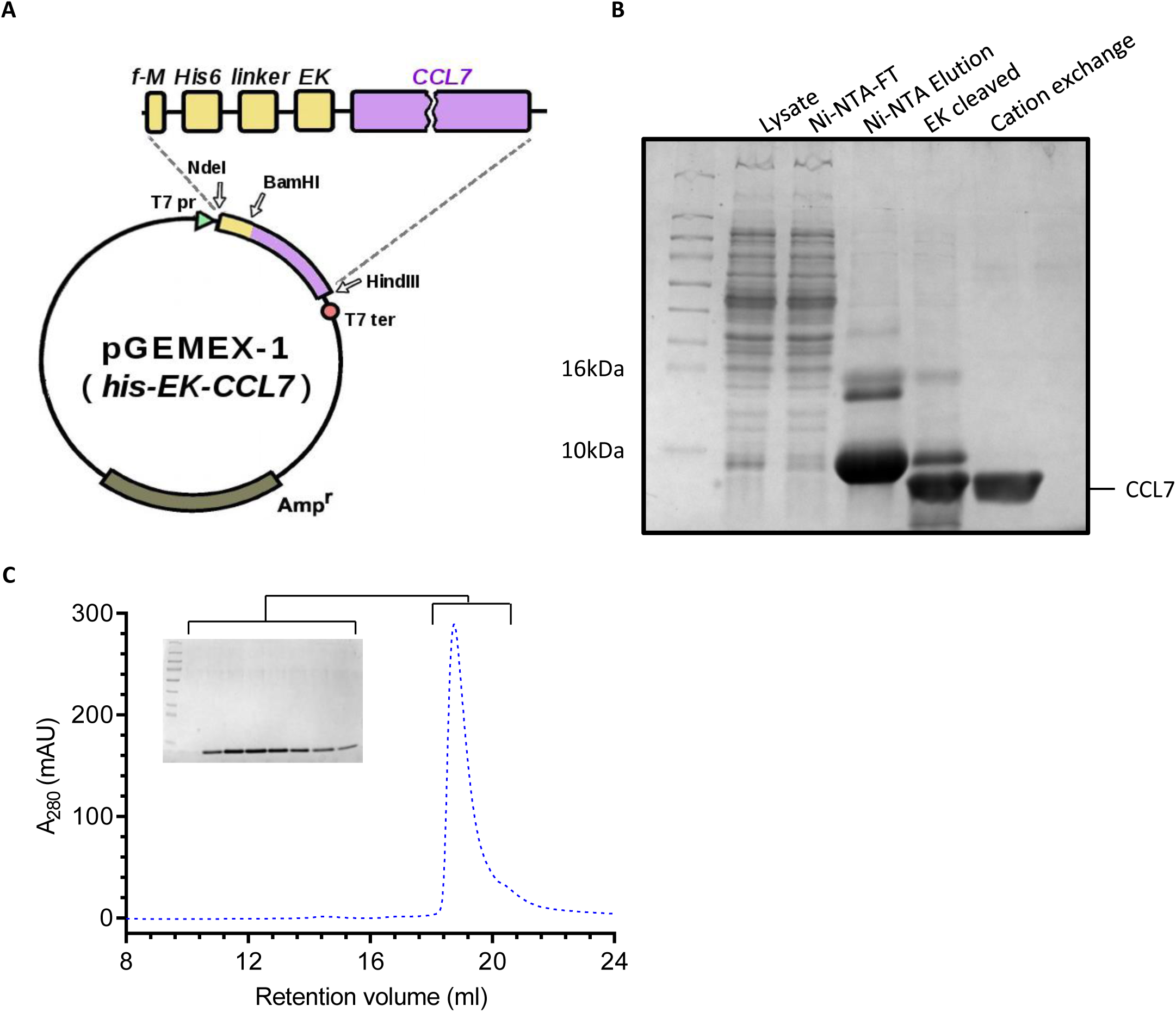
Expression and purification of native CCL7 from *E. coli*. **A.** pGEMEX-1 derived expression construct for recombinant production of native CCL7. The natural signal sequence of CCL7 was removed during cloning of CCL7 and an N-terminal His6 tagged followed by enterokinase (EK) cleavage site (DDDDR) were engineered. A five amino acid long linker (GSGSG) was also included between the his-tag and EK site. **B.** Purification of CCL7 using Ni-NTA, enterokinase digestion, and cation exchange chromatography yielded purified protein. Samples from every step were loaded and separated using SDS-PAGE followed by Coomassie staining for visualization. **C.** Size exclusion chromatography of CCL7 on Superdex 200 column revealed a monodisperse peak with an elution volume of 20mL. The inset show the corresponding fractions analyzed by SDS-PAGE and Coomassie staining.

### Functional characterization of native CCL7 in G-protein coupling and β-arrestin recruitment assays

We measured the functionality of purified native CCL7 using two different assays. First, we carried out GloSensor assay as described above to evaluate the ability of native CCL7 to induce Gαi-coupling for CCR2. Unlike his-CCL7, the native CCL7 was as potent and efficient as CCL7-Fc with an IC_50_ of about 0.3nM (Figure 5A-B). These observations suggest that the his6-tag present at the N-terminus of CCL7 potentially interferes with its ability to induce effective G-protein-coupling at CCR2. Second, we carried out confocal microscopy based assay to assess whether native CCL7 can effectively drive βarr recruitment for two different 7TMRs namely the ACKR2 and CCR2. As presented in Figure 6A-D, we observed efficient membrane translocation of βarr2 at early time-points (2-5 min) and subsequent translocation to endosomes upon extended ligand stimulation (5-30 min), for both CCR2 and ACKR2. These data demonstrate that native CCL7 purified from *E. coli* is functional in terms of inducing efficient receptor-transducer coupling, and therefore, suitable for structure-function studies in future.

**Figure 5:**
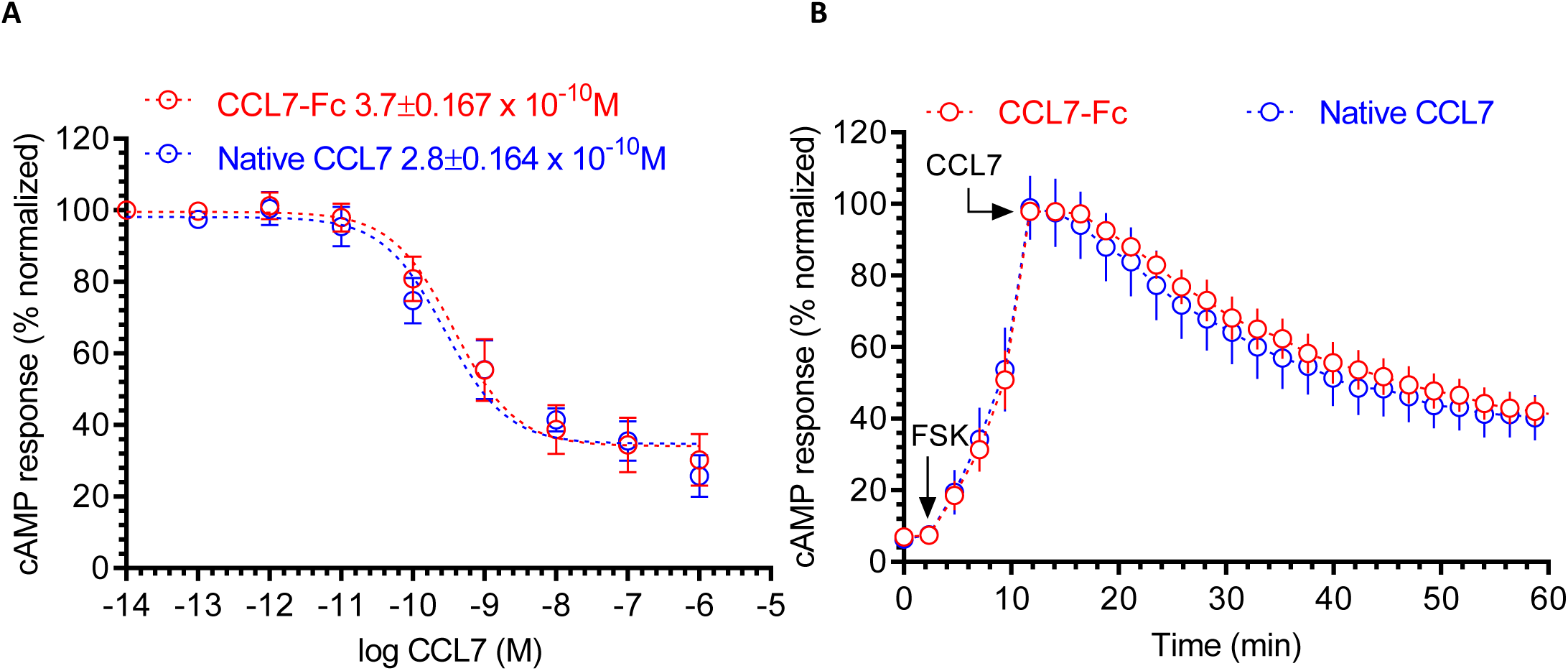
Functional characterization of native CCL7 in GloSensor based G-protein coupling assay. **(A)** The ability of different CCL7 preparations to inhibit Forskolin-induced cAMP response downstream of human chemokine receptor CCR2 in transfected HEK-293 cells. Native CCL7 represents untagged version while CCL7-Fc represents a C-terminal Fc-tagged version of CCL7 expressed and purified from *Sf*9 cells. Data represent average±SEM of four independent experiments carried out in duplicates and normalized with respect to lowest concentration of CCL7-Fc (treated as 100%). (B) Time-course analysis s of CCL7 induced decrease in the cAMP level over the indicated time-period. Values recorded in the GloSensor assay at a concentration of 1nM from the experiments presented in panel A are plotted. The arrow indicates the time of CCL7 addition and the values are normalized with maximal cAMP response observed for CCL7-Fc (treated as 100%).

**Figure 6:**
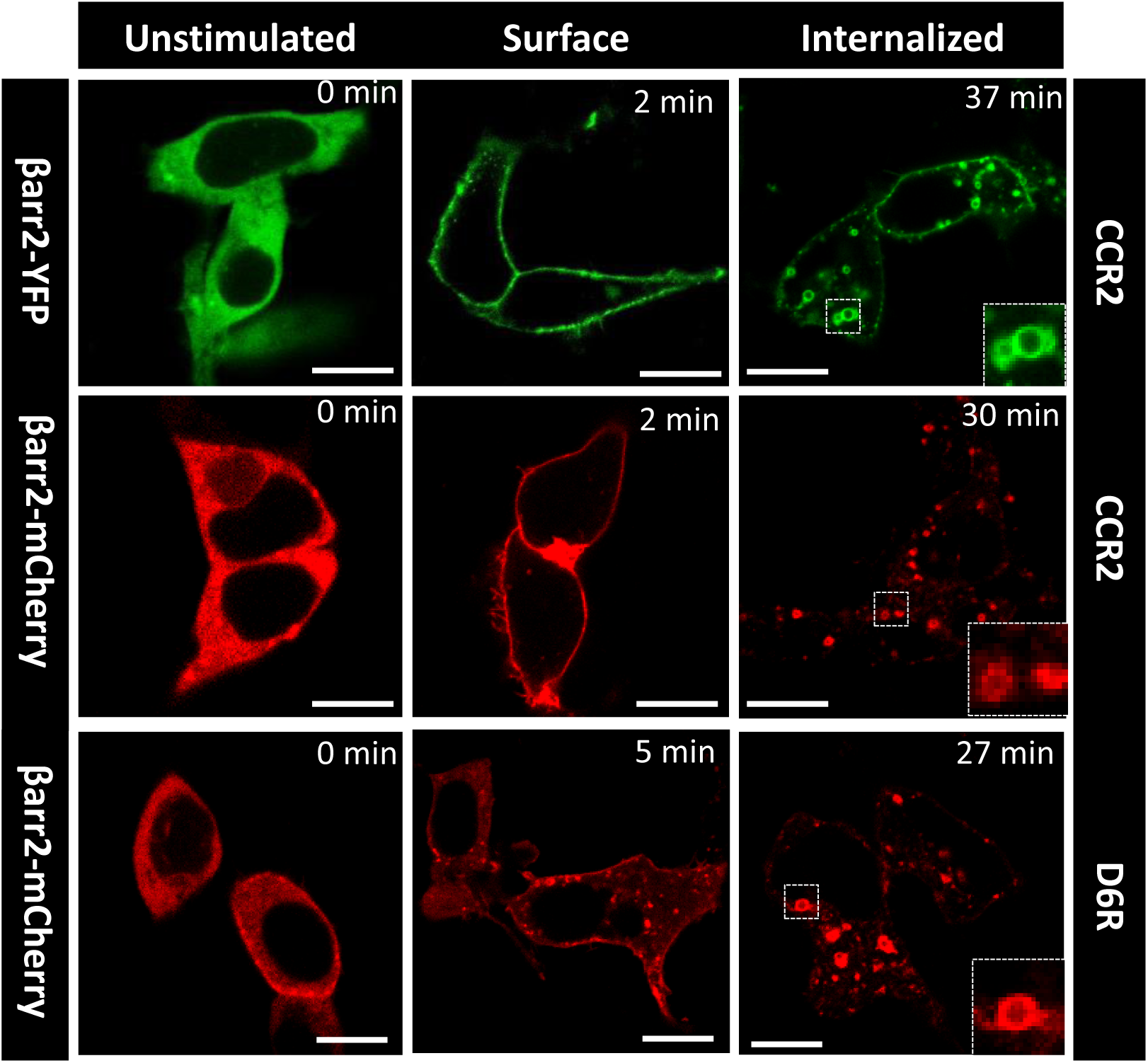
Functional characterization of CCL7 as assessed by agonist-induced trafficking of β-arrestin 2. HEK-293 cells expressing either CCR2 or D6R (ACKR2) together with β-arrestin 2-YFP or β-arrestin 2-mCherry were stimulated with saturating concentration of native CCL7 (1μM) for indicated time-points. Subsequently, the trafficking of β-arrestin 2 was monitored using confocal microscopy. Representative images from two independent experiments are shown and the scale bar is 10μm.

## Discussion

Investigating the interaction of chemokines with chemokine receptors is an important subject area in order to understand their activation, signaling and regulatory mechanisms. However, technical challenges associated with preparing large amounts of chemokines in native and fully functional form has limited our ability to study structural aspects of ligand-receptor complexes for the members of chemokine receptor sub-family. Although some functional studies have used chemokines in *E. coli*, they have primarily used constructs that do not yield native N- and C-terminus resulting in sub-optimal affinity and efficacy, and making them less relevant for structural characterization. Furthermore, in most cases, chemokines expressed in *E. coli* were present in inclusion bodies and had to be refolded for subsequent studies; however, our protocol yielded soluble expression of CCL7 that can be purified in milligram amounts without the need for any refolding procedure.

Our strategy based on the incorporation of enterokinase cleavage site at the N-terminus of CCL7 in the fusion protein allows us to preserve native sequence without any residual modification arising from affinity tags or cleavage sites. This results in expected potency profile of CCL7 in Gαi-coupling for CCR2 and in promoting typical trafficking pattern of βarrs for ACKR2 and CCR2. Comparison of cAMP response induced by his-CCL7 and native-CCL7 also indicates that modification of the N-terminus of CCL7 compromises its functionality with respect to transducer coupling, an observation that may be relevant to other chemokines as well. It is plausible that a similar strategy can be adapted for generating the milligram amounts of native version of other chemokines as well as their isotope-labeled derivatives, and it should facilitate structure-function studies of chemokine receptors.

An obvious caveat of expression of chemokines in *E. coli* is the lack of post-translational modifications (PTMs) such as glycosylation. Although there is evidence of O-glycosylation for one of the chemokines, CCL2, when isolated from native tissues, it appears to have no significant effect on activity (Jiang et al., 1991, Jiang et al., 1990, Proost et al., 2006). Most of the chemokines appear to be functional even in the absence of any PTM, however, this is an aspect that should be investigated further with respect to receptor binding and transducer coupling. Our experiments suggest that native CCL7 purified from *E. coli* is fully functional with respect to Gαi-coupling for CCR2 and βarr recruitment for CCR2 and ACKR2. Nonetheless, future studies evaluating signaling outputs and cellular responses may be important to further establish the complete functional profile of CCL7.

In conclusion, we present a streamlined strategy for preparing milligram amounts of fully functional CCL7 with native N- and C-terminus, which can potentially be adapted for other chemokines. Our study paves the way for structural and functional characterization of CCL7-bound CCR2 and ACKR2, and potentially other chemokine receptors.

## Acknowledgments

The work described in this paper is primarily supported by an Indo-Russian project [INT/RUS/RFBR/P-253] from the Department of Science and Technology [DST] awarded to AKS in India and Russian Foundation for Basic Research (RFBR) grant 17-54-45064 in Russia awarded to ASA. In addition, the research program in Dr. Shukla’s laboratory is supported by an Intermediate Fellowship of the Wellcome Trust/DBT India Alliance Fellowship [grant number IA/I/14/1/501285] awarded to AKS, the Science and Engineering Research Board (SERB) (EMR/2017/003804), Innovative Young Biotechnologist Award from the Department of Biotechnology (DBT) (BT/08/IYBA/2014-3) and the Indian Institute of Technology, Kanpur. Dr. Shukla is an Intermediate Fellow of Wellcome Trust/DBT India Alliance, EMBO Young Investigator and Joy Gill Chair Professor. Drs. Hemlata Dwivedi and Mithu Baidya were supported by National Post-Doctoral Fellowship of SERB (PDF/2016/002930 and PDF/2016/2893). Dr. Ashish Srivastava is supported by the Wellcome Trust/DBT India Alliance Early Career Fellowship [grant number IA/E/17/1/503687]. The sequential NMR assignment of ^15^N-labeled his-CCL7 and secondary structure verification in Prof. Arseniev’s laboratory are supported by Russian Science Foundation (RSF) grant 19-74-30014 awarded to ASA. We thank Dr. Eshan Ghosh and Ravi Ranjan for their contribution in purification of CCL7-Fc from *Sf*9 cells, Igor A. Ivanov for mass spectrum of his-CCL7.

## Author contributions

MVG designed and executed the purification of his-CCL7 and native CCL7, produced isotope-labeled variant of his-CCL7 for NMR experiments; MAD and KDN performed NMR experiments and analyzed NMR data; DR carried out expression and purification of CCL7 in the Shukla laboratory with assistance from AS, and participated in GloSensor and confocal microscopy experiments executed by HD and MB, respectively. All authors contributed in writing and editing the manuscript. AKS and ASA managed the overall project.

## Materials and methods

### General reagents, strains and cell lines

XL-1 and BL21(DE3)pLysS strains of *E.coli* were purchased from Stratagene (USA) and a previously described vector pGEMEX-1(his6) was used for generating expression constructs (Goncharuk et al., 2018). Synthetic oligonucleotides were produced and DNA sequencing was performed by Evrogen (Russia). Isotope labels (^2^H, ^15^N, ^13^C) were incorporated using CIL reactives (United States).

### Construction of the expression plasmids

The human CCL7 gene was PCR amplified without the intrinsic signal sequence along with partial codon optimization for expression in *E. coli* from the pFastBac-CCL7_Fc vector (designed and generated in Shukla lab) using specific primers. For his-EK-CCL7, a sequence encoding DDDDR amino acids (i.e. enterokinase cleavage site) was introduced at the 5’ end of the CCL7 gene. The PCR products were cloned into home-made pGEMEX-1(his6) vector (Goncharuk et al., 2018) using NdeI and HindIII restriction sites. Expression plasmids were verified by DNA sequencing.

### Expression of CCL7 in *E. coli*

Expression plasmids encoding his-CCL7 and his-EK-CCL7 were transformed in chemically competent BL21(DE3)pLysS cells and plated onto LB or YT agar plates supplemented with 100μg/ml ampicillin and 25 mkg/ml chloramphenicol. After overnight incubation at 37°C, 50 medium-size single colonies were flushed with 1ml TB media to inoculate 1L TB media along with 200μg/ml ampicillin and 25 mkg/ml chloramphenicol. Bacterial cells were cultured at 27°C, 250 rpm overnight until next day when the OD_600_ reached 1.5. At this point, culture was induced with 1mM IPTG, and the cultures were grown at 20°C for additional 48h. Subsequently, cells were harvested using centrifugation and pellets were store at −80°C.

In order to prepare isotope labeled CCL7, transformed cells were cultured using M9 minimal salt medium, containing 0.0002% yeast extract and (0.2% ^15^NH_4_Cl), (0.2% ^15^NH_4_Cl, 0.4% [U-^13^C]-glucose), or (D_2_O, 0.2% ^15^NH_4_Cl, 0.4% [U-^13^C]-glucose), correspondingly. Cell were induced at an OD600 of 0.6 with 0.5mM IPTG followed by subsequent growth at 27°C for additional 48h.

### Purification of his-CCL7 and native CCL7

Cell pellets corresponding to 1L culture were re-suspended in 50ml lysis buffer containing 30mM MOPS, pH 7.2, 1M NaCl, 10mM imidazole, 5%(v/v) glycerol, 0.3%(v/v) TritonX-100, 1mM PMSF. Cells were lysed by sonication on ice for 15 min with amplitude of 45 % (pulse of 15sec ON and 30sec OFF) or until complete lysis took place and cell lysate was then centrifuged at 14000g for 1h at 4°C followed by filtration through 0.22μm filter to remove cellular debris and unbroken cells. Clear supernatant was loaded onto a Ni-NTA affinity column containing 4ml Ni-NTA resin (Clontech) pre-equilibrated with lysis buffer at an approximate flow rate of 1ml/min. Flow through fraction containing unbound proteins was collected and the column was washed with 25 bed volumes of wash buffer containing 30mM MOPS, pH 7.2, 1M NaCl, 10mM imidazole, and 5%(v/v)glycerol. Bound proteins were then eluted with 6-7 bed volumes of elution buffer (30mM MOPS, pH 7.2, 1M NaCl, 500mM imidazole, and 5% glycerol). The elution fractions were analyzed on SDS-PAGE and fractions containing maximum amount of CCL7 were pooled and used for subsequent second purification step.

For his-CCL7, Ni-NTA eluate was desalted by about ten-fold dilution with 30 mM MES buffer, pH 5.8 or by dialysis against the 30 mM MES, 30 mM NaCl buffer, pH 5.8, clarified by centrifugation at 24000g for 20min at 4°C, filtered through a 0.22 μm filter and loaded onto to a cation-exchange column (5mL, SP FF, GE Healthcare), pre-equilibrated with 30 mM MES, pH 5.8, 30mM NaCl. Afterwards, the column was washed until the UV (280nm) reading reached baseline, and subsequently, bound proteins were eluted with a 30-1000 mM linear NaCl gradient in ten column volumes. Peak fractions were analyzed by SDS-PAGE, pooled and concentrated/desalted using 3kDa Amicon spin-concentrators, flash frozen and stored at −80°C in small aliquots for subsequent experiments. For NMR experiments, isotope-labeled CCL7 were buffer exchanged to 5mM Potassium Phosphate buffer, pH 5.1 containing, 0.0025% NaN_3_.

For the purification of native CCL7, Ni-NTA elution of his-EK-CCL7 was dialyzed overnight at 4°C against enterokinase digestion buffer (20mM Tris, pH 8.0, 50mM NaCl, 2mM CaCl_2_). Precipitated proteins were removed by centrifugation at 14000g for 30 min at 4°C and the remaining soluble protein was incubated with enterokinase light chain (NEB) for 16h at room-temperature as per manufacturer’s protocol. The cleavage efficiency of enterokinase under these conditions was typically 70-80%. Afterwards, native CCL7 was isolated using two different and independent protocols. In the first method, enterokinase cleaved sample was loaded on to Resource S cation exchange column followed by elution using a linear gradient of NaCl (50-1000mM) over 10 column volumes. In the second method, enterokinase cleaved sample was loaded onto 1mL Ni-NTA resin and flow through fractions containing native CCL7 were collected. In both methods, fractions containing native CCL7 were pooled and stored as described above. Expression and purification of CCL7-Fc construct using baculovirus infected *Sf*9 cells will be described separately (Manuscript in preparation).

### Size-exclusion chromatography

Purified his-CCL7 and native CCL7 were loaded on pre-equilibrated (20mM Hepes, 100mM NaCl, pH 7.5) Superdex 200 Increase (10/300 GL) column (GE) in a volume of 100μl at 1-10mg/ml concentration. Subsequently, the column was run at 0.4 ml/min flow rate, the elution profile of CCL7 was monitored at 280nm and fractions were analyzed on SimplyBlue stained SDS-PAGE.

### NMR spectroscopy

NMR spectra of “cold” and ^15^N-labeled his-CCL7 were acquired on Bruker Avance 700MHz NMR spectrometer equipped with 5 mm PATXI probe. The following NMR spectra were acquired in H_2_O/D_2_O (9:1), pH 5.1 (pH-meter readings), temperature 30°C: 2D NOESY (τ_m_=100ms) and 2D TOCSY (τ_m_=60ms) for “cold” his-CCL7 sample; 2D ^1^H-^15^N HSQC, 3D ^1^H-^15^N NOESY-HSQC (τ_m_=100ms) and 3D ^1^H-^15^N TOCSY-HSQC were acquired for ^15^N-labeled his-CCL7, 3D spectra were acquired and processed in Non-Uniformly Sampled mode (Kazimierczuk and Orekhov, 2015).

### GloSensor assay

The ability of purified CCL7 to trigger G-protein coupling was assessed with respect to inhibition of forskolin-induced cAMP response using GloSensor assay as described previously (Pandey et al., 2019, Kumari et al., 2017, Kumar et al., 2017). Briefly, HEK cells at a density of 3 million were transfected with 3.5μg each of CCR2 and the luciferase-based cAMP biosensor (pGloSensorTM-22F plasmid; Promega) plasmids. After 16-18 hour of transfection, cells were trypsinised, centrifuged and resuspended in buffer (1XHBSS and 20mM HEPES, pH 7.4) containing 0.5mg/ml luciferin (LUCNA-1G/GOLDBIO). The cells were then seeded in a 96 well plate at a density of 6 × 10^4^ cells /100μl/ well. The plate was kept at 37°C for 90min in the CO_2_ incubator followed by incubation at room temperature for 30 minutes. Basal reading was read on luminescence mode of multi-plate reader (Victor X4). Since CCR2 is a Gαi-coupled receptor, a receptor-independent adenylyl cyclase stimulator, Forskolin (10μM) was added to each well and luminescence was recorded until reading were stable (10 cycle repeats). Thereafter, cells were stimulated with varying doses of each ligand ranging from 0.01pM to 1μM and luminescence was recorded for 60min using a microplate reader. Data was normalized with maximal response obtained with highest ligand concentration after basal correction and analyzed using nonlinear regression in GraphPad Prism software. Data presented in Figure 3 and 5 were measured simultaneously with shared CCL7-Fc condition.

### Confocal microscopy

In order to test the functionality of CCL7 in terms of βarr recruitment and trafficking, we used confocal microscopy based analysis of βarr2 recruitment and trafficking as described previously (Ghosh et al., 2017, Pandey et al., 2019). Briefly, HEK-293 cells were transfected using either ACKR2 or CCR2 (3.5 μg each per 10cm plate) along with either βarr2-YFP or βarr2-mCherry (3.5 μg). 24h post-transfection, cells were seeded at one million density on a 35×10 mm confocal dish (GenetiX). To study agonist dependent βarr2 recruitment in transfected cells, live cell imaging was carried out 48 h post-transfection using the Zeiss LSM 710 NLO confocal microscope with oil-immersion 63X /1.40 NA, objective, equipped with CO_2_ and temperature controlled platform and having 32x array GaAsP descanned detector (Zeiss). A multiline argon laser at 488 nm and a Diode Pump Solid State Laser at 561 nm was used for imaging YFP-tagged βarr2 or mCherry-tagged βarr2 respectively. CCL7 (1μM) was added to the cells and incubated for ∼2 mins and the cells were then imaged for up to 40 min.

### Quantification and statistical analysis

Expression and purification experiments were repeated multiple times, and functional assays were repeated at least three times. Corresponding details of data normalization and quantification are included in the respective figure legends.

